# Identifying conserved molecular targets required for cell migration of glioblastoma cancer stem cells

**DOI:** 10.1101/669036

**Authors:** Josephine Volovetz, Artem D. Berezovsky, Tyler Alban, Yujun Chen, George F. Aranjuez, Ashley Burtscher, Kelly Shibuya, Daniel J. Silver, John Peterson, Danny Manor, Jocelyn A. McDonald, Justin D. Lathia

## Abstract

Glioblastoma (GBM) is the most prevalent primary malignant brain tumor and is associated with extensive tumor cell infiltration into the adjacent brain parenchyma. However, there are limited targeted therapies that address this disease hallmark. While the invasive capacity of self-renewing cancer stem cells (CSCs) and their non-CSC progeny has been investigated, the mode(s) of migration used by CSCs during invasion is currently unknown. Here we used time-lapse microscopy to evaluate the migratory behavior of CSCs, with a focus on identifying key regulators of migration. A head-to-head migration assay demonstrated that CSCs are more invasive than non-CSCs. Time-lapse live cell imaging further revealed that GBM patient-derived CSC models either migrate in a collective manner or in a single cell fashion. To uncover conserved molecular regulators responsible for collective cell invasion, we utilized the genetically tractable *Drosophila* border cell collective migration model. Candidates for functional studies were generated using results from a targeted *Drosophila* genetic screen followed by gene expression analysis of the human homologs in GBM tumors and associated GBM patient prognosis. This strategy identified the highly conserved small GTPase, Rap1a, as a potential regulator of cell invasion. Alteration of Rap1a activity impaired the forward progress of *Drosophila* border cells during development. Rap1a expression was elevated in GBM and associated with higher tumor grade. Functionally, the levels of activated Rap1a impacted CSC migration speed out of spheres onto extracellular matrix. The data presented here demonstrate that CSCs are more invasive than non-CSCs, are capable of both collective and single cell migration, and express conserved genes that are required for migration and invasion. Using this integrated approach, we identified a new role for Rap1a in the migration of GBM CSCs.

## Introduction

Glioblastoma (GBM), the most prevalent primary malignant brain tumor, remains one of the most lethal cancers, with a median survival of less than two years despite aggressive interventions including surgery, radiation, and chemotherapy^1, 2, 3, 4^. Barriers to effective treatment of GBM include the extensive infiltration of tumor cells throughout the brain and the high degree of inter- and intra-tumoral heterogeneity; these properties have long been recognized as histological hallmarks of GBM. Another hallmark of GBM is the self-renewing cancer stem cell (CSC) population that is resistant to conventional therapies^5, 6, 7^. While many studies have identified mechanisms through which CSCs expand and are resistant to radiation and Temozolomide, the standard of care chemotherapy, there is less known about the molecular mechanisms that drive invasion. Several studies have reported that CSCs display elevated invasive potential over non-CSCs^8, 9, 10^. However, the direct differences between CSC and non-CSC invasion are unclear, as previous assessments have been done by comparing cell populations in isolation. A number of molecular pathways that drive GBM cell invasion, such as the Wingless/Int1 (Wnt) and TGF-β pathways^11^, have been identified, with efforts underway to neutralize these pathways as potential therapeutic interventions. Since these mechanisms are also associated with self-renewal^12^, the extent to which they drive invasion over self-renewal has yet to be determined. Increased invasion has been observed with anti-angiogenic therapies^13, 14^. Additionally, GBM cells visualized *in vivo* tend to migrate along the luminal surface of blood vessels; the perivascular environment thus allows the cells to migrate faster and induces further microvascular development^15, 16^. These observations suggest that invasion itself may contribute to therapeutic resistance in GBM.

The mode by which GBM cells, particularly CSCs, migrate and invade the brain is poorly understood. Studies using tumor histology, live *ex vivo* tumor explants, and *in vivo* models demonstrate that cancer cells have the capacity to invade as single cells or collectives, in which cells coordinate their movement by maintaining cell-cell contacts amongst small to large groups of cells^17, 18^. Both single cell and collective cell modes of invasion have been observed in a variety of human tumors, including breast, thyroid, and lung carcinomas^19, 20, 21, 22, 23, 24^. Additionally, migrating cancer cells are highly dynamic and can invade as single cells, loosely-associated multicellular streams, collective nests or strands, or through expansive growth, with some cells changing their migration mode during movement in response to the surrounding tissue microenvironment^17, 25^. Recent work has demonstrated that GBM cells can migrate as both single cells and as multicellular collectives, which may influence their capacity to infiltrate the surrounding brain parenchyma^26, 27^. However, whether GBM CSCs themselves migrate and invade as cell collectives, and whether this differs from non-CSCs, has yet to be determined and is the focus of our studies.

Although collective cell invasion contributes to cancer, much of our current mechanistic understanding of how cells migrate as collective groups have been obtained by studying cells that move during normal development. Indeed, collective cell migration is a frequent mode of cell migration in the embryo, where it contributes to the shaping and forming of many organs. Key examples include gastrulation to form the three embryonic germ layers, neural crest migration to give rise to craniofacial structures and the peripheral nervous system, zebrafish lateral line organ formation, and branching morphogenesis to elaborate tubular structures within organs (e.g. the mammary gland and mammalian lung)^28, 29^. One of the best-studied models of developmental collective cell migration is the *Drosophila* border cells, which migrate as a cohesive group of six to ten cells in the egg chamber, the functional unit of the ovary^29^. The border cell cluster migrates during oogenesis in two phases, both of which respond to specific ligands secreted by the oocyte: in the posterior phase, border cells undergo a long-range movement from the anterior end of the egg chamber to the oocyte at the posterior; in the dorsal phase, the cells undergo short-range migration along the oocyte towards the dorsal-anterior side of the egg chamber^29, 30^. The ability to genetically manipulate and observe border cell migration in its native tissue environment in real time makes it a powerful tool for identifying conserved regulators of collective invasion in development and in cancer^29, 31, 32^. Moreover, the use of the *Drosophila* system has also recently been leveraged for studies to identify conserved molecular mechanisms that drive GBM cell proliferation, survival, and self-renewal^33, 34, 35, 36, 37^.

In this study, we identified patient-derived GBM CSC models that migrate as mixtures of collectives and individual cells, as well as several lines that primarily move as collective strands. Further, we used results from a *Drosophila* border cell screen to identify conserved genes that control cell migration, which represent potential targetable regulators of GBM CSC invasion. Through this approach, we identified Rap1a as one of the downstream effectors of this screen. We found that human Rap1a levels were elevated in GBM, and altered Rap1a activity impacted live CSC migration. Taken together, these data demonstrate the ability to identify molecular regulators of migration and invasion of GBM CSCs, including Rap1a, using a multi-system approach.

## Results

### CSCs are more invasive than non-CSCs

Previous studies have suggested that CSCs have increased migration and invasion capacity compared to non-CSCs. However, these analyses were done on separate populations of CSCs or non-CSCs, and not together in a competition assay that would normalize for confounding factors such as media conditions or paracrine- and autocrine-secreted factors. To assess whether there were functional differences in cell invasion, we compared differentially-labeled CSCs and non-CSCs in a head-to-head co-culture ECM-based assay (**Fig. 1A**). We used an approach previously shown to accurately assess co-culture invasion in breast cancer^38^. We fluorescently labeled CSCs and non-CSCs, then seeded them, overlaid the cells with a 3D extracellular matrix, and then added a chemoattractant on top. Using this system, we compared three patient-derived GBM CSC models, T387, T4121, and T3691, versus their corresponding non-CSC progeny, all of which were independently derived from patient-derived xenograft (PDX) models. After 24 hours, we imaged the cells on a spinning disk confocal to assess the extent of invasion into the matrix along the chemokine gradient. With this head-to-head migration assay, differentially-labelled CSCs versus non-CSCs were evaluated directly in the same dish for their migratory capacity. In all models, we observed significantly more invasion by CSCs compared to non-CSCs (**Fig. 1B-C**). Overall, CSCs exhibited 2- to 5-fold increase in migration over non-CSCs. These results thus demonstrate that CSCs are more invasive than non-CSCs when compared in identical conditions.

**Figure 1.**
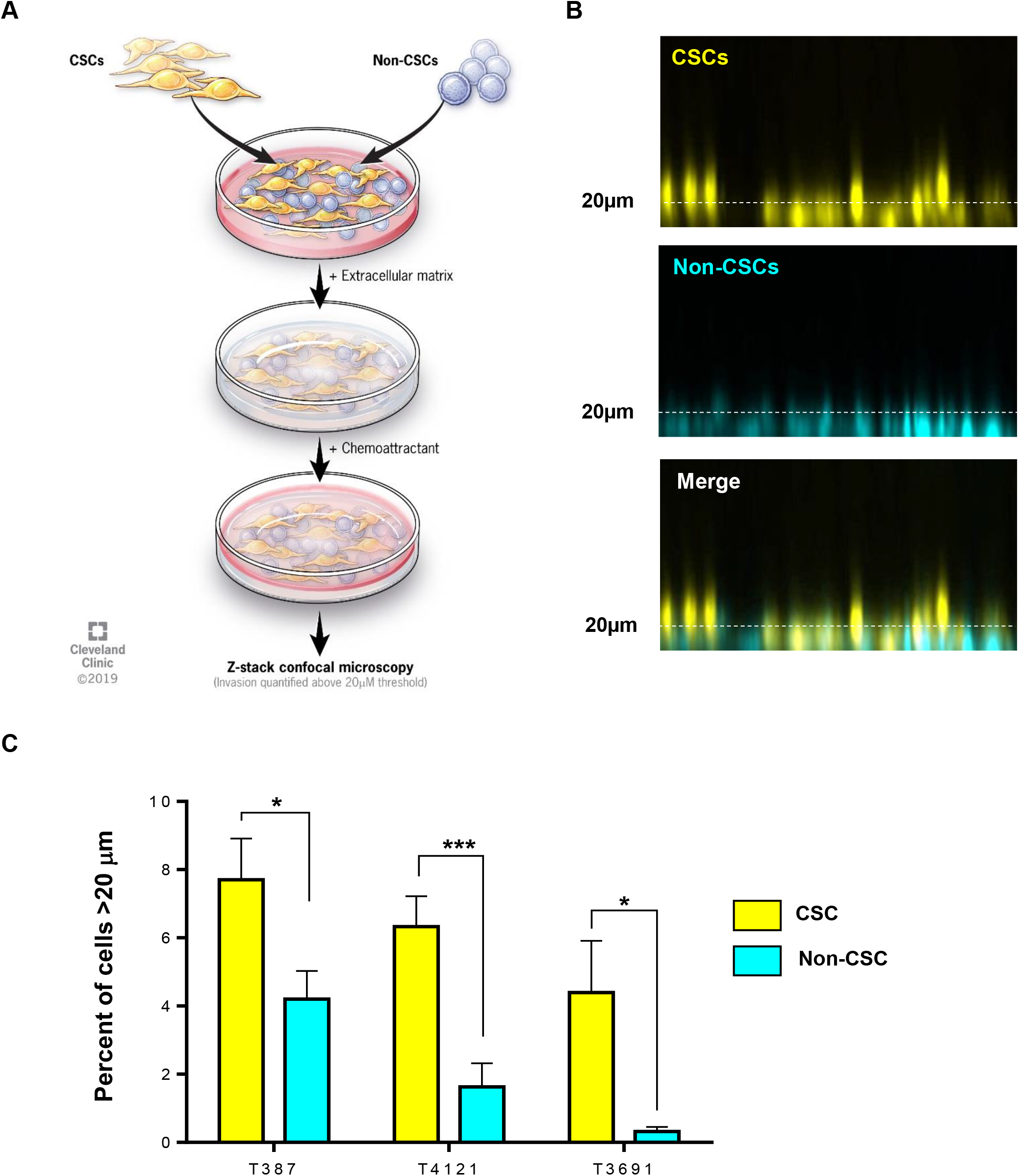
Head to head migration of cancer stem cell and non- cancer stem cells. Schematic representation of the head-to-head migration assay of cancer stem cells (CSCs) and non-CSCs embedded into a 3D Geltrex extracellular matrix with a chemoattractant layered on top (**A**). Representative confocal *z*-stack projections of CSCs and non-CSCs 12 hours post-seeding (CSCs = yellow, non-CSCs = blue, **B**). Extent of migration of CSCs and non-CSCs from the bottom of the well (**C**). Statistics calculated based on an unpaired student’s t-test, *p<0.05, ***p<0.001.

### Different CSC lines display different modes of migration

To further assess how CSCs invade in terms of single cell and collective cell migration, we next analyzed how CSCs exit sphere culture when plated on extracellular matrix. In this assay, eight separate CSC models were grown as spheres before being introduced to an extracellular matrix coated surface and then imaged using time-lapse microscopy to capture the movement of cells out of the sphere and onto the matrix. All CSCs were able to migrate away from the original sphere. Some models migrated individually (T4121 and T3832), whereas others migrated as cell collectives, most often as cohesive “finger-like” projections or small nests (T387 and L1; **Fig. 2** **and Supplemental Fig. 1**). In the collectively migrating lines, the cells consistently stayed connected, with N-cadherin enrichment at contacts between cells (**Fig. 2**; **Supplemental Fig. 2**). The majority of CSC models, however, displayed a mixture of migration modes, with cells moving away from the tumor sphere both as individuals and as collectives, most often in small cell groups (T3691, GBM10, L0, and T1919; **Fig 2**, **Supplemental Fig. 1**). These data together demonstrate that CSCs employ different modes of cell invasion, highlighting yet another phenotype that displays inter-tumor and intra-tumor heterogeneity.

**Figure 2.**
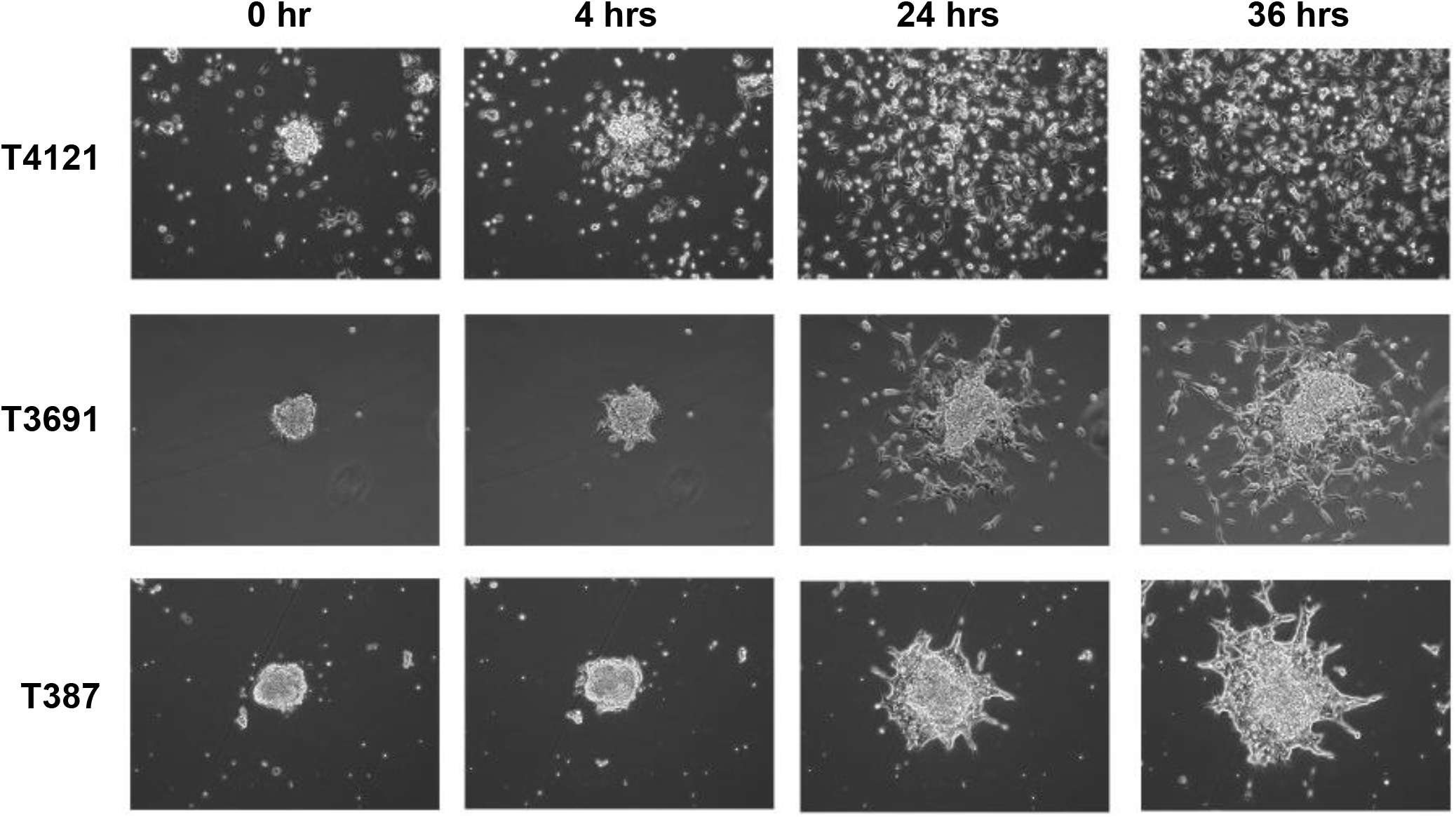
Modes of migration utilized by GBM cancer stem cells. Time-lapse microscopy of cells from three patient-derived glioblastoma (GBM) CSC models (T4121, T3691, T387) exiting from a sphere after 0, 4, 24, and 36 hours (hrs) in culture.

### Identification of candidate CSC invasion genes via border cell migration

Given the observation that CSCs display different modes of cell invasion, the majority as collectives or as mixtures of single cells and collectives, we sought to determine the underlying mechanisms driving CSC migration patterns. *Drosophila* border cells represent a genetically tractable model of collective cell migration within an intact tissue. Many genes known to regulate border cell migration are highly conserved in humans and have been implicated in cancer^31, 39, 40^. During mid-oogenesis, six to ten epithelial follicle cells are recruited to form the cohesive border cell cluster, which migrates as a coordinated unit over the course of about four hours towards the oocyte located at the posterior of the egg chamber^41^.

Recently, we performed an RNA interference (RNAi) screen targeting PDZ domain-containing genes to identify those that regulate border cell collective migration^42^, which provided a starting point to identify candidate GBM migration genes. Proteins that contain the PDZ protein-protein interaction domain facilitate the formation of multi-protein scaffolding complexes with conserved roles in signaling, cell polarization, and adhesion, making them excellent candidates to regulate the collective migration of normal and cancer cells^43, 44^. The majority of *Drosophila* PDZ-(<u>P</u>SD95/<u>D</u>lg/<u>Z</u>O1-) domain genes were screened for the ability to promote border cell migration^42^ (**Fig. 3A**). This screen identified high confidence PDZ-domain-containing genes (multiple transgenic RNAi lines targeting the gene were able to disrupt migration; **Supplemental Table 1:** Group 1 genes) and lower confidence genes (only one RNAi line per gene disrupted migration; **Supplemental Table 1**: Group 2 genes). We further included well-characterized genes known to promote border cell migration and/or interact with PDZ domain genes (**Supplemental Table 1**: Group 3 genes)^30, 39^.

**Figure 3.**
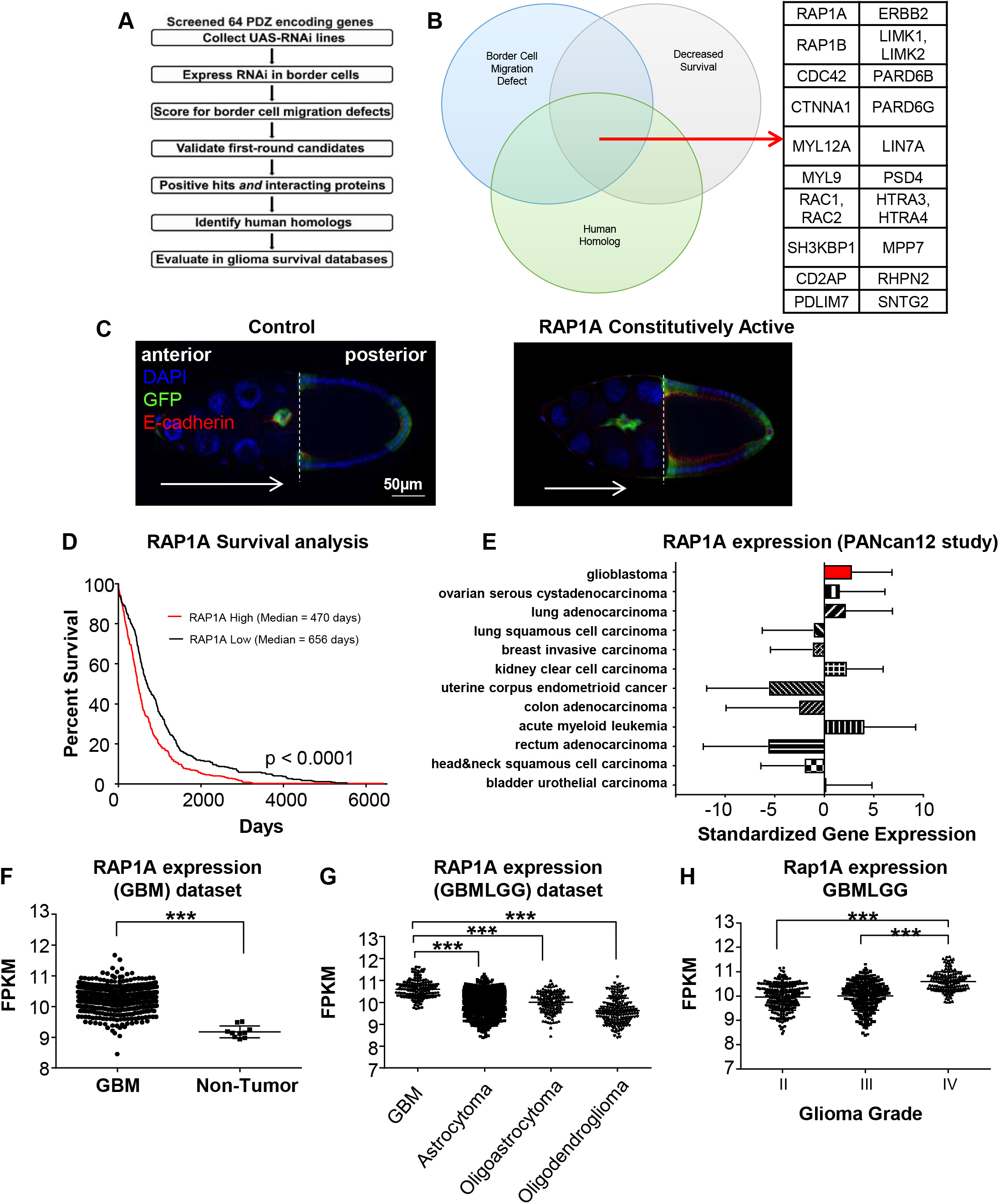
Identification of Rap1a as a potential GBM invasion regulator. Flowchart of approach to identify human homologs of fly genes that are required for migration and correlate with human glioma patient prognosis (**A**). Venn diagram of *Drosophila* genes involved in border cell migration, with annotated human homologs, and genes associated with decreased survival of GBM patients (**B**). Rap1a is one of twenty-three genes that meet these criteria. Control border cells (left panel, **C**) in the *Drosophila* egg chamber collectively migrate towards the oocyte (dashed line; extent and direction of migration indicated by white arrow). Expression of constitutively active Rap1a (right panel, **C**) results in failure to reach the oocyte and altered cluster shape. Border cells express UAS-mCD8:GFP (green) driven by *slbo*-GAL4. Egg chambers were immunostained for E-cadherin (red), which is highly expressed in border cells and other cell membranes. Nuclei were visualized using DAPI (blue). Evaluation of human Rap1a expression in the cancer genome atlas GBMLGG database and its association with glioma survival (**D**). Rap1a expression across the pan cancer database of tumor mRNA expression demonstrates increased Rap1a in GBM and some other cancer types (**E**). Analysis of Rap1a expression in GBM versus non-tumor identified increased Rap1a expression in GBM tumors compared to non-tumors (**F**). Analysis of Rap1a expression in GBMLGG dataset identified increased expression in GBM tumors compared to other glioma tumors (**G**). Analysis of Rap1a expression in GBMLGG dataset identified increased Rap1a expression with increasing glioma grade (**H**). FPKM: fragments per kilobase of transcript per million mapped reads. Statistics calculated based on one-way ANOVA, ***p<0.001 and differences in survival calculated based on log-rank analysis.

We identified human orthologs for these 40 *Drosophila* migration genes using FlyBase and the Drosophila RNAi Screening Center Integrative Ortholog Prediction Tool (DIOPT; http://www.flyrnai.org/diopt)^45^. The identified human orthologs were then used to interrogate the TCGA-GBMLGG data set in order to find overlapping genes that were relevant in both the migration model and glioma patients. Genes whose high expression correlated with decreased patient survival were compared to the list of genes that decreased border cell migration (**Fig. 3B**; **Supplemental Table 1**). Twenty-three genes fit these criteria and were candidates for further functional tests in border cells and GBM CSCs. Notably, genes encoding small GTPases, or targets of small GTPases, were well-represented (**Fig. 3B**). Small GTPases are known to promote both single cell and collective cell migration, and many interact with PDZ-domain proteins^46, 47, 48^. For example, the high-confidence hit LIMK1 is downstream of Rac (human Rac1 and Rac2), which promotes membrane protrusions in migrating cells^49, 50^. Similarly, Cdc42 stimulates actin-rich protrusions and can polarize migrating cells through direct binding to Par-6 (human PARD6B/PARD6G), one of the strongest hits from the border cell screen^51, 52, 53^, and has a well-described role in GBM invasion^54^.

As a validation of the identified candidate genes (**Fig. 3A, B**; **Supplemental Table 1**), we confirmed that the small GTPase Cdc42 was required for migration in multiple models. In border cells, expression of a dominant-negative (DN) mutant form of Cdc42 (Cdc42^N17^; DN Cdc42) severely blocked migration, with most border cells stopping along the migration pathway and failing to reach the oocyte (**Supplemental Fig. 3A-C**). Live border cells expressing DN Cd242 overall had trouble initiating migration compared to controls (**Supplemental Fig. 3C**; **Supplemental Movies 1, 2**). Further, Cdc42 controlled migratory protrusions (**Supplemental Fig. 3D**). We found that DN Cdc42 border cell clusters extended more protrusions, and these protrusions were more persistent, with a longer lifetime than those produced by control border cells. These data are consistent with a recent study that found that Cdc42 promotes cell-cell communication amongst border cells, resulting in only one border cell at the front of the cluster being able to extend a productive protrusion, which facilitates migration of the entire collective to the oocyte^55^. Increasing the levels of Cdc42 via overexpression of wild-type (WT) Cdc42 also modestly impaired movement to the oocyte, although to a lesser extent than DN Cdc42 (**Supplemental Fig. 3A, C**). These results suggest that optimal levels of the Cdc42 small GTPase are required for migration *in vivo*.

Similarly, in GBM, we reduced the activity of Cdc42 in T387 cells using a specific chemical inhibitor, ML 141^56^. Decreased Cdc42 reduced the cell migration out of a CSC sphere (**Supplemental Fig. 4A**), but not cell viability, survival or proliferation (**Supplemental Fig. 4B-D**). Our results are consistent with recent work by Okura et al.^54^, which found that knockdown of Cdc42 by siRNA decreased GBM invasion. Further, constitutively active Cdc42 reduced survival in a PDX model of glioma, while increasing the invasive capacity in sphere exit assays^54^. Thus, high Cdc42 correlates with GBM progression through increased invasion.

### Rap1a levels are elevated in GBM patients and correlate with tumor grade

Having established that this integrated approach can identify genes important for GBM CSC migration, we next focused on a candidate gene, human Rap1a, which is less characterized in collective cell migration. We and others recently showed that inhibition of the *Drosophila* Rap1a homolog (Rap1) disrupted migration to the oocyte due to defects in actin-rich protrusions and altered cell-cell adhesion^57, 58^. Rap1a is also regulated by the screen multi-hit gene, PDZ-GEF^42^ (ortholog of human RapGEF2/PDZ-GEF1 and RapGEF6/PDZ-GEF2; Supplementary Table 1). Further, expression of constitutively active (CA) Rap1a impaired movement of the border cells to the oocyte (**Fig. 3C**)^57, 58^. Thus, similar to Cdc42, having the proper levels of Rap1a is critical for successful collective migration of border cells. In GBM, Rap1a expression was found to be increased compared to non-tumor tissue using the TCGA database, and increased expression correlated with increasing tumor grade (**Fig. 3 F-H**). Additionally, when compared across the pan-cancer dataset, GBM had elevated levels of Rap1a compared with other common cancers, although it was not ubiquitously increased across all cancers (**Fig. 3E**). Survival analysis in the GBMLGG dataset also revealed that Rap1a expression levels correlate with a poor prognosis (**Fig. 3D**).

### Activated Rap1a alters migration of CSCs

On the basis that Rap1a levels are generally elevated in GBM, and that *Drosophila* Rap1 is required for collective migration of border cells (**Fig. 3A-B**)^57^, we wanted to determine how altering Rap1a levels and/or activity impacted CSC migration. As described above, T387 CSCs normally migrate as collective fingers and small clusters (**Fig. 2**). In comparison with wild-type Rap1a, constitutively-active Rap1a (63E) did not alter the migration rate (**Fig. 4A-C**). Both conditions had reduced migration when compared to an untransfected control. However, dominant-negative (N17) Rap1a induced a slight increase in migration rate compared to other overexpression conditions and had a similar migration rate to control conditions (**Fig. 4A-C**). These data suggest that levels of active Rap1a are critical for CSC migration.

**Figure 4.**
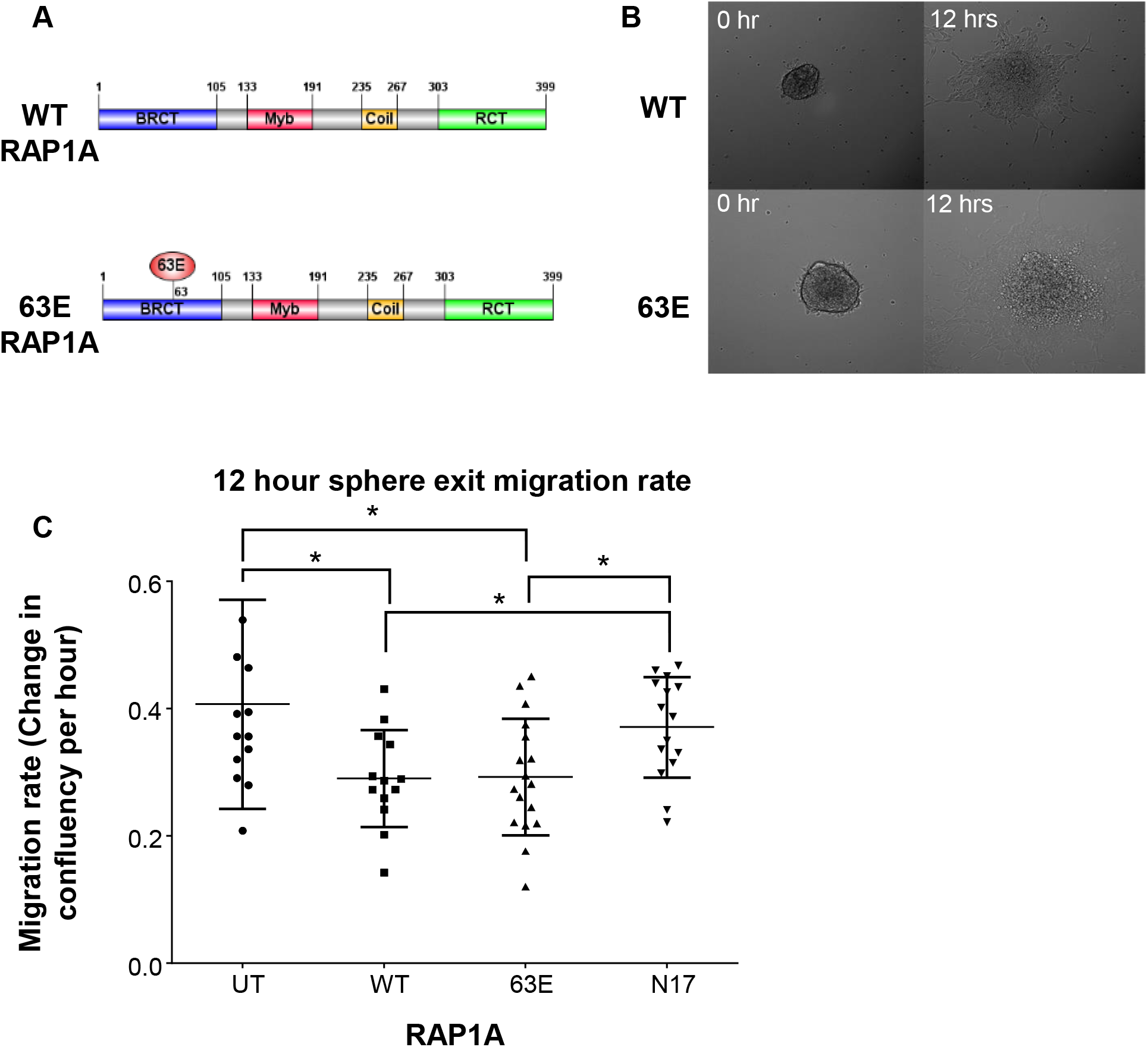
Rap1a function in GBM CSCs. 3D extracellular matrix spheroid time-lapse migration assay of patient-derived GBM CSC model (T387) transfected with wild-type (WT) or constitutively active (63E) Rap1a (**A**). Stills from time-lapse microscopy of GBM CSC cells transfected with WT or 63E Rap1a migrating from a sphere at 0 hr and 13 hrs (**B**). Quantification of migration rate between Rap1A constructs (WT, 63E, N17) and untransfected (UT) control cells (**C**). Statistics calculated based on a student’s t test or one-way ANOVA, *p<0.05.

## Discussion

Cancer cells can migrate individually or as collectives^18, 19, 20, 29, 59^. The mode of migration may be critical for how well tumor cells, including those in GBM, are able to invade the surrounding tissue and disseminate. Several recent studies found that GBM cells can migrate collectively^26, 27^, although whether GBM CSCs specifically use this type of migration is an open question. Our data indicate that a variety of patient-derived GBM CSC models can move in small to large collectives away from tumor spheres plated on Geltrex, although CSCs can also migrate individually and as mixtures of single cells and collectives. Interestingly, previous studies indicated that CSCs are likely more invasive than non-CSCs in GBM^8, 9, 10^. However, these analyses were done separately, so it was unknown whether CSCs could outcompete non-CSCs during invasion. In this study, we directly compared the invasive capacity of CSCs to non-CSCs, by co-incubating CSCs and non-CSCs in the same dish, then allowing both cell types to migrate through an ECM matrix together over time. Our results with this head-to-head Geltrex-based assay allowed for greater control and comparison, and definitively showed that CSCs are more invasive than non-CSCs. Currently it is unknown whether the mode of CSC invasion directly contributes to progression of GBM in patients. Recent work by the Friedl group demonstrated that human glioma cells can invade as multicellular groups, as well as individual cells, into 3D astrocyte scaffolds, mouse brain slice cultures, mouse brain xenografts, and in human tumor samples^26, 27^. Specifically, glioma cells maintain cell-cell connections while moving along both blood vessels and the astrocyte-rich brain stroma^26^. Thus, we favor a model in which collective invasion of GBM CSCs helps disseminate tumors into the brain parenchyma and vasculature^60^, although this has yet to be directly tested.

Because GBM invasion is a dynamic and multi-step process, the molecular mechanisms are still poorly understood. The results presented here highlight unique opportunities to apply knowledge about migration and invasion from model organisms to human diseases such as GBM. Border cell migration during *Drosophila* oogenesis has become a valuable tool for studying collective cell migration due to the wealth of genetic and cell biological methods that allow for wild-type and mutant cells to be imaged *ex vivo* while migrating within the native tissue^61^. We used an informed approach to identify genes required for border cell migration that may have a role in GBM. In particular, we focused on the highly conserved PDZ-domain encoding genes, many of which have known or suspected roles in human tumorigenesis and cancer cell invasion^44, 62, 63^. Of these PDZ genes, plus related genes such as small GTPases with validated functions in border cells^30^, we identified a number of conserved genes whose elevated expression correlated with decreased GBM patient survival. The results presented here, as well as recent results by others, indicate that the small GTPases Cdc42 and Rap1A are needed for the migration of both border cells and GBM CSCs^54, 55, 57, 58^. Further work will be needed to determine the roles, if any, of the additional identified genes on GBM tumor growth, migration, and/or invasion. Given the high conservation of human disease genes in *Drosophila*, especially those implicated in tumor invasion and metastasis^31^, our results further suggest that border cell migration can be used to identify additional relevant regulators of GBM cancer cell migration and invasion.

Rap1a GTPase is known to regulate the migration of both normal cells and cancer cells^64, 65^. Rap1 belongs to the Ras family of small GTPases, but is regulated by distinct guanine nucleotide exchange factors (GEFs) and GTPase activating proteins (GAPs), and generally has Ras-independent functions^65^. Rap1a (*Drosophila* Rap1) was recently found to be required for collective migration of *Drosophila* border cells, specifically in the outer migratory border cells of the cluster where Rap1 promotes the formation of protrusions that help keep the border cells motile^57, 58^. Interestingly, Rap1 in border cells also keeps the cells in the border cell cluster tightly adhered together through cell-cell adhesions^58^. In this context, both decreased and elevated Rap1a activity impaired the migration of border cells, by altering the polarity and adhesion of cells within the collective as well as border cell movement upon their migratory substrate, the nurse cells. Similarly, here we showed that altered levels of human Rap1a impacted the migration speed of GBM CSCs. Accumulating evidence indicates a general role for Rap1a in the cell migration and invasion of a variety of tumors^64, 66^. Depending on the cancer, either elevated *or* decreased Rap1a activity enhances or suppresses tumor invasion, typically through regulation of either cadherin- or integrin-based adhesions^66^. This suggests complex roles for Rap1a and the downstream molecular pathways and effectors, which likely depend on the tumor type and cellular context^64^.

Here we found that constitutively-active Rap1a decreased the migration speed of CSCs compared to dominant-negative Rap1a. However, elevated Rap1a levels correlate with worse GBM patient outcome. Additionally, Rap1 increases U87MG glioma spheroid invasion on collagen in response to platelet-derived growth factor (PDGF) stimulation^67^. One way to reconcile these counterintuitive findings is to consider the possibility that Rap1a may have different roles in GBM CSCs versus non-CSCs, as well as unknown effects on tumor progression *in vivo* versus cell culture. Moreover, it is also possible that CSCs have an optimized level of Rap1a activity, and our overexpression conditions compromised this equilibrium. We hypothesize that speed and migration persistence could impact how tumors with elevated Rap1a expression invade the brain. Although here we transiently increased Rap1a expression using a wild-type Rap1a construct and similarly observed slower migration, this still may not reflect what happens in GBM tumors in the native environment. Chronic elevation of Rap1a expression in GBM tumors could be worse for disease progression than transient activation. Interestingly, Rap1a has been shown to be important for cellular proliferation in the context of *in vivo* GBM tumor growth^68^. It will be important in future to test whether tumors that overexpress Rap1a are more infiltrative *in vivo*, whether the main function of Rap1a in GBM is in cell proliferation^68^, or whether some combination of tumor growth, migration, and invasion is at play. The effect that Rap1a has on GBM patient survival is likely to be complex, and not simply one in which increased Rap1a expression increases tumor invasion and thus decreases patient survival.

An open question is what are the molecular and cellular mechanisms by which altered levels of Rap1a impact GBM CSC migration, invasion, and patient survival. Gain of Rap1a function slowed the migration of CSCs compared to dominant-negative Rap1a. One possibility is that Rap1a may regulate the adhesion of CSCs to their migratory ECM substrate, with elevated levels of Rap1a increasing adhesion and slowing down migration. Similarly, integrins help cells migrate by adhesion to ECM^69^. In particular, optimal migration speed occurs with intermediate levels of integrins and adhesion strengths, while slower migration occurs with either low or high levels of integrins^69, 70^. Given that Rap1 is required for activation of some integrins (e.g. alpha-II beta-3 integrin) and regulates integrin-based adhesions in a variety of cell types^71, 72, 73^, future work should focus on whether Rap1a regulates GBM CSC adhesion to ECM through integrins. In some contexts, activation of Rap1a is dependent upon junctional adhesion molecule-A (JAM-A), Afadin, and PDZ-GEF2, all of which are known to associate with each other^74^. Dimerization of JAM-A is necessary for epithelial cell migration and leads to protein complex formation with Afadin and PDZ-GEF2^74, 75^. This in turn is associated with an increase in the activation of Rap1a and cell migration. Although these studies were performed in epithelial cells, the relationship between JAM-A and Rap1a is intriguing due to our previous observations that JAM-A is necessary and sufficient for GBM CSC maintenance^76, 77^. Thus, in the future, it will be important to explore any relationship between JAM-A and Rap1a in GBM CSCs and the potential impacts this would have on GBM invasiveness and tumor growth. Taken together, our data provide a rationale for additional studies on the role of Rap1a in GBM migration and invasion. These studies based on reducing cell migration and invasion should be a high priority due to the limited availability of treatments in this category, which remains difficult to target and is a driver of recurrence. Finally, our approach highlights the ability to leverage model organisms to identify key processes that drive GBM invasion and highlight a paradigm that could be applied to many other key cancer processes.

## Materials and Methods

### Isolation of CSCs

CSCs were isolated from surgically obtained GBM specimens and transiently amplified by implantation of unsorted tumor cells into immunocompromised mice. Derived xenografts were dissociated before culture. All specimens were generated under approved Institutional Review Board protocols. For xenograft dissociation, single cells were prepared from the bulk tumor by a Papain dissociation kit (Worthington Biochemical) as per the manufacturer’s protocol and cultured using previously reported culturing methods^76, 77^. Cells were maintained in serum-free growth media supplemented with EGF and bFGF (“neurobasal complete media”) as spheres or adherent on Geltrex solution (Thermo Fisher Scientific).

### Head-to-head invasion assay

The head-to-head invasion assay was adapted from Goswami et al.^38^ with minor modifications. First, cancer cells were stained with either Vybrant DiO or DiI cell labeling solutions (Thermo Fisher Scientific), according to manufacturer’s directions. 150,000 to 200,000 cells in neurobasal complete media, from two distinct cell types, were added 1:1 to 35 mm glass bottom dishes (MatTek Corporation) coated with a dilute layer of Geltrex matrix. After 24 hours, media was removed and a thick, 80 μl layer of Geltrex matrix was layered over the adherent cells. Dishes were returned to the incubator for 30 minutes, in order for the matrix to solidify. Then, the dishes were flooded with 3 mL of neurobasal complete media. The following day, live cells were visualized using a PerkinElmer UltraView VOX Spinning Disk Confocal Imaging System on a Nikon DM1-6000 SD inverted microscope using a 10×/NA 0.7 air objective. A 100-μm z-stack image at 5-μm increments was obtained. An extended *x*–*z* projection was then obtained using Volocity software (PerkinElmer) and invasion was quantified using ImageJ (NIH, Bethesda, MD, USA) as the percentage of Vybrant dye signal found above 20 μm from the bottom of the dish, compared with total fluorescence within the field.

### Immunofluorescence

For immunofluorescence analysis of adherent cultures, cells were fixed with 4% paraformaldehyde (Millipore Sigma, Burlington, MA,USA) at room temperature for 15 minutes, washed three times with phosphate-buffered saline (PBS) and blocked with 5% normal goat serum (Millipore) in PBS plus 0.1% Triton X-100 for 1 hour. Cells were incubated with an anti-N-cadherin (1:200; Cell Signaling) antibody. Cells were washed three times with PBS plus 0.1% Triton X-100 (Millipore) and incubated with appropriate secondary antibody (1:100 anti-mouse IgG3 Alexa 488 and 1:300 goat anti rabbit Alexa 568; Thermo Fisher Scientific). Nuclei were counterstained with DAPI solution, a coverslip added, and mounted on slides using VECTASHIELD Hardset mounting media (Vector Labs). Fluorescence micrographs were acquired using a Leica SP5 confocal microscope, and images were processed in Adobe Photoshop CS6 (Adobe).

The *Drosophila* strain *slbo*-GAL4, UAS-mCD8:GFP was crossed to *w*^*1118*^ (control) or the following UAS-transgene flies to drive expression in border cells: UAS-Rap1^V12^ (constitutively-active Rap1), UAS-Cdc42^N17^ (dominant-negative Cdc42), UAS-Cdc42 (wild-type Cdc42). Detailed information about *Drosophila* strains can be found in FlyBase (http://flybase.org/). Ovaries from the correct progeny of these crosses were dissected, fixed in 4% methanol-free formaldehyde (Polysciences) in phosphate buffer, and stained for anti-E-cadherin (DCAD2, Developmental Studies Hybridoma Bank (DSHB), University of Iowa) at 1:10 dilution using standard protocols^78^. Secondary antibodies used were anti-mouse or anti-rat Alexa 568 at 1:400 (Thermo Fisher Scientific). Nuclei were labeled using 0.05 μg/ml DAPI (Millipore). Ovaries were further dissected and mounted on slides in Aqua-PolyMount (Polysciences), followed by imaging on a Zeiss AxioImager Z1 microscope. Images were taken using a Zeiss Axiocam 503 camera, the Zeiss ApoTome2 module, and a 20 × 0.75 NA objective, using either Zeiss AxioVision 4.8 or Zen software.

### Time-lapse microscopy of border cell migration in *ex vivo* cultured egg chambers

Live imaging was performed essentially as previously described^79, 80^. Briefly, Drosophila egg chambers were dissected from whole ovaries and cultured in Schneider’s insect medium (Thermo Fisher Scientific) supplemented with 10% fetal bovine serum (Millipore Sigma), 0.20 mg/mL insulin (Millipore), and penicillin/streptomycin (Thermo Fisher Scientific). Egg chambers were transferred onto a 50 mm Lumox® gas-permeable bottom dish (Sarstedt, Inc.), overlaid with a 22 × 22 mm coverslip and the edges sealed with halocarbon oil 27 (Millipore Sigma) to prevent evaporation.

Widefield fluorescence *z*-stack images (5 *z*-sections at 3μm apart) were captured every 2 minutes for the duration of border cell migration (about 3 to 4 hours). The Zeiss Colibri light source (25% blue LED at 250 msec exposure, full frame ROI) was used to illuminate GFP, which was expressed in border cells using the GAL4-UAS system. Movies were created using the Zeiss Axiovision 4.8 “Inside 4D” software module or the ImageJ distribution Fiji (http://fiji.sc)^81^. For each time-lapse movie, concurrent *z*-sections in which the border cell cluster was in focus were merged. Image brightness and/or contrast were adjusted in Fiji.

Protrusions were defined manually, as in Sawant et al.^57^. Essentially, the converted QuickTime movies were analyzed using Fiji. Cellular extensions that were longer than 10 pixels (at least about 4 μm) from the main cell body were considered to be “protrusions.” Only those protrusions that appeared within the first hour of each movie were tracked for analyses. Statistical tests were performed in GraphPad Prism 7.

### Time-lapse microscopy of tumor spheres

Floating spheres were transferred into Geltrex-coated 6-well dishes. Live time lapse movies of adherent CSC cells were acquired using a Leica DMI6000 inverted microscope and LAS X software v3.4.18368.2 (Leica Microsystems) equipped with a Hamamatsu ImageEM CCD camera and a Hamamatsu Orca Flash4 camera (Hamamatsu Photonics). Phase contrast images of multiple fields per well were collected every 15 minutes for 36 hours using a 10x 0.4 NA objective lens. Image processing and analysis was performed using the open source analysis program Fiji (64 bit, build 2014.11.25)^81^. Change in confluency of CSC cells over time was quantified for 3 different time points from each of the time lapse movies as a measure of cell proliferation and migration. Selected phase contrast images were pre-processed by first converting them to 32 bit images and then using the Contrast Limited Adaptive Histogram Equalization (CLAHE) plugin to increase contrast and flatten image background. Image segmentation and quantification of confluency of the CLAHE processed phase contrast images was then completed using the PHANTAST plugin^82^.

### Cell manipulations

Cancer stem cells (CSCs) were transiently transfected in 10 cm plates. Once cells achieved a confluency of about 75%, they were transfected with a preparation of OptiMEM, 14.37μg of DNA, and FuGENE HD Transfection reagent (Promega). On day 2 after the transfection, cells were removed from their plates with Accutase (BioLegend) and counted. For Rap1a alterations, CSCs were transfected with RAP1 WT, RAP1 N17, RAP1 63E, and with an eGFP control. Cdc42 was inhibited in T387 cells using ML 141 (Sigma) at 200 μM. Rap1 WT and N17 constructs were obtained from previously published sources^83^. Rap1 63E was obtained from Addgene (Plasmid #32698) from previously published sources^83, 84^. The eGFP was obtained from Lonza.

### Cell viability assays

2000 cells were added to Geltrex-coated 96-well plates. Opaque clear bottom plates were used for both CellTiterGlo and CaspaseGlo (Promega) assays and regular cell culture plates were used for CyQUANT assay (Thermo Fisher Scientific). Cells were incubated at 37°C and 5% CO2 for the duration of the experiment. Assays were completed according to manufacturer’s directions on 1- and 3-days post-plating or addition of the inhibitor. CaspaseGlo measurements were normalized to CellTiterGlo measurements.

### Statistics

All statistical analyses were performed in GraphPad Prism 5.0 (GraphPad Software). Unless otherwise indicated, a student’s t test or one-way ANOVA was used and a *p* value of ≤0.05 was considered significant.

## Acknowledgments

This work used the Leica SP5 confocal/multi-photon microscope that was purchased with partial funding from National Institutes of Health SIG grant 1S10RR026820-01. This work was supported in part by a fellowship from the Kansas INBRE through the National Institutes of Health grant P20 GM103418 to Y.C., NIH R21 CA198254 to J.D.L. and J.A.M., and support from the Cleveland Clinic (including the Brain Tumor Research and Therapeutic Development Center of Excellence) and Case Comprehensive Cancer Center to J.D.L.

