## SupplementalFigures for "Identifying conserved molecular targets required for cell migration of glioblastoma cancer stem cells"

**Supplemental Figure 1. GBM CSC lines exhibit a variety of migration modes.** Time-lapse microscopy of cells from patient-derived GBM CSC models (T3832, GBM10, L0, T1919, L1) exiting from a sphere over 36 hours (hrs).

**Supplemental Figure 2. N-cadherin and localization in T387 tumorspheres.** Confocal micrographs of CSCs (T387) migrating as a collective out of a tumorsphere stained with an antibody against N-cadherin (red, left panel). Closer view of peripheral migrating group (in box) is shown (right panel). Nuclei stained with 4′,6-diamidino-2-phenylindole (DAPI, blue). Scale bars represent 50μm and 25μm as indicated on individual micrographs.

**Supplemental Figure 3. Cdc42 levels correlate with GBM patient survival and border cell migration.** Cdc42 requirement during border cell collective migration (**A-D**). Control border cells (arrow) are shown at different stages of migration (top row), from early stage 9 (e9) at the start of migration up to stage 10 when they reach the oocyte. Expression of wildtype (WT) or dominant-negative (DN) Cdc42 in border cells (bottom row) using *slbo*-GAL4 both result in migration defects, as shown by the failure to reach the oocyte by stage 10. *slbo*-GAL4 was used to drive UAS-mCD8:GFP (green) expression in border cells; E-cadherin (red) labels all cell membranes; nuclei were visualized by DAPI (blue). Scale bar represents 20 µm (**A**). Stills from time-lapse movies of control border cells (top row) or border cells expressing DN Cdc42 ($Cdc42^{DN}$, bottom row) during migration. Control border cells start their movement, migrate, and reach the anterior side of the oocyte (o; arrowheads). Border cells expressing $Cdc42^{DN}$ fail to migrate. Border cell clusters (arrows) are visualized by *slbo*-GAL4-driven expression of mcD8:GFP (**B**). Quantification of the frequency of various migration outcomes of control border cells (WT, black bars) or border cells overexpressing wild-type (WT) Cdc42 (red bars) or DN Cdc42 (blue bars) in stage 10 egg chambers. Error bars represent standard deviation; n $\geq$ 50 egg chambers in each of 3 trials (**C**).

Quantification of protrusion number (left) and lifetime (right panel) in control (n = 8 movies) and

$Cdc42^{DN}$ (n = 10 movies) border cell clusters (**D**). All data points are shown; whiskers represent the minimum and maximum measurements, the box extends from the 25th to 75th percentiles, and the line indicates the median. Statistics calculated based on an unpaired two-tailed t test (**C**) or one-way ANOVA (**D**), *p<0.05, **p<0.01, ***p<0.001.

**Supplemental Figure 4. Requirement for Cdc42 in GBM CSC migration.** Representative micrographs of CSC-enriched spheres from a patient-derived xenograft (L1) model treated with a Cdc42 inhibitor (ML141, 200 μM) demonstrates limited migration as compared with a DMSO control (**A**). Quantification of cell viability (**B**), survival (**C**), and proliferation (**D**) of CSC-enriched spheres treated with a Cdc42 inhibitor (ML141, 200 μM) demonstrates no major changes with inhibitor treatment versus control. Values represent means +/- standard deviation (n = 4).

**Movie 1. Time-lapse video of control border cell migration.** Representative example of a live stage 9 control (*slbo*-GAL4, UAS-mCD8:GFP) egg chamber. Frames from this video are shown in Supplemental Figure 3C (top panels). Border cells expressing GFP form a cluster and delaminate from the epithelial follicle cells, then migrate between the unlabeled nurse cells as a cohesive group, finally reaching the unlabeled oocyte at the posterior (time in minutes). Frames were acquired every 2 min with a 20x objective. Anterior is to the left.

**Movie 2. Time-lapse video of DN Cdc42 border cell migration.** Representative example of a live stage 9 DN Cdc42 (*slbo*-GAL4, UAS-mCD8:GFP/UAS-$Cdc42^{N17}$) egg chamber. Frames from this video are shown in Supplemental Figure 3C (bottom panels). DN Cdc42 border cells expressing GFP extend extra protrusions, take longer to delaminate, and do not reach the oocyte at the posterior. Frames were acquired every 2 minutes with a 20x objective. Anterior is to the left.

**Supplemental Figure 1**

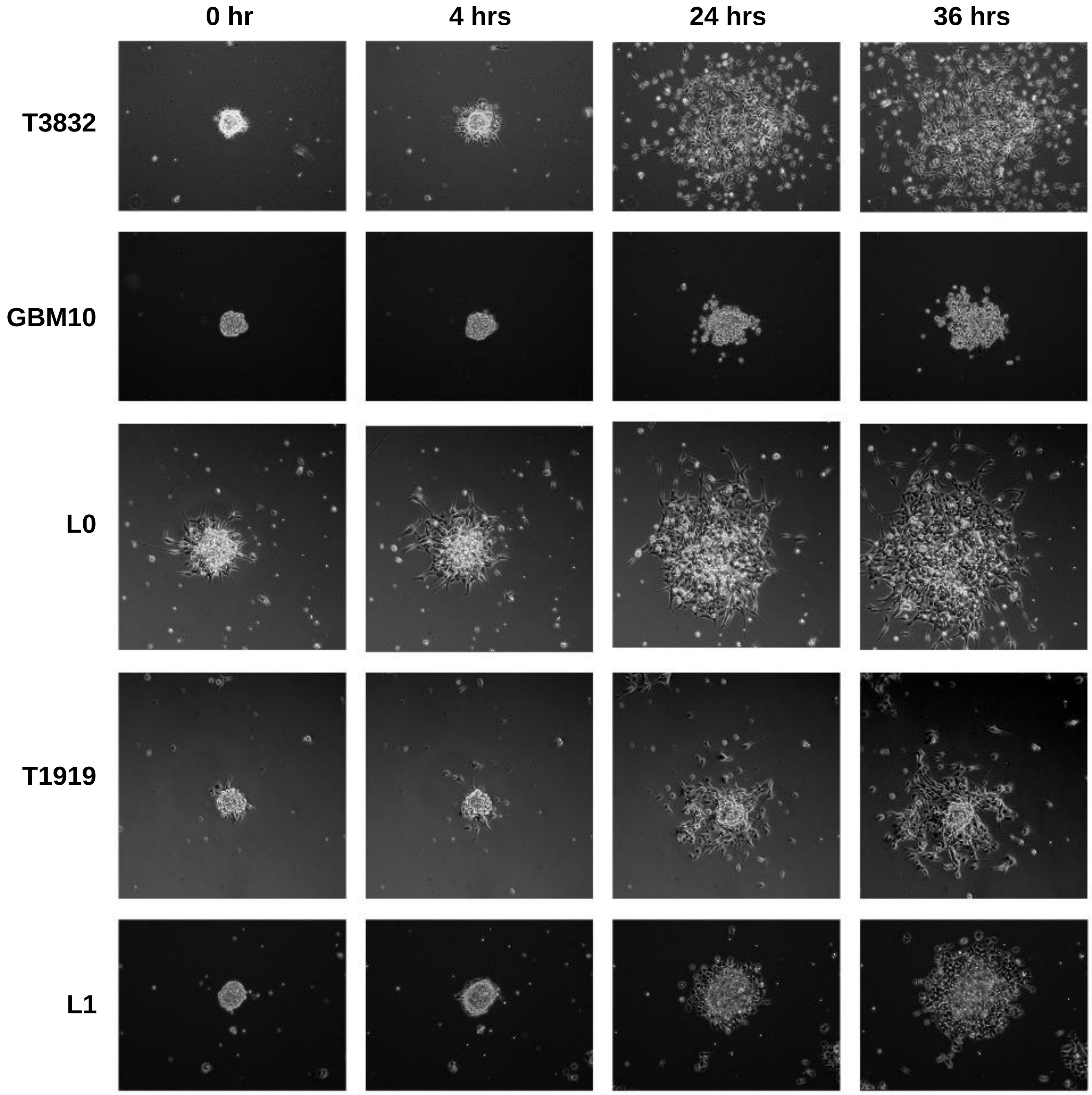

**Supplemental Figure 2**

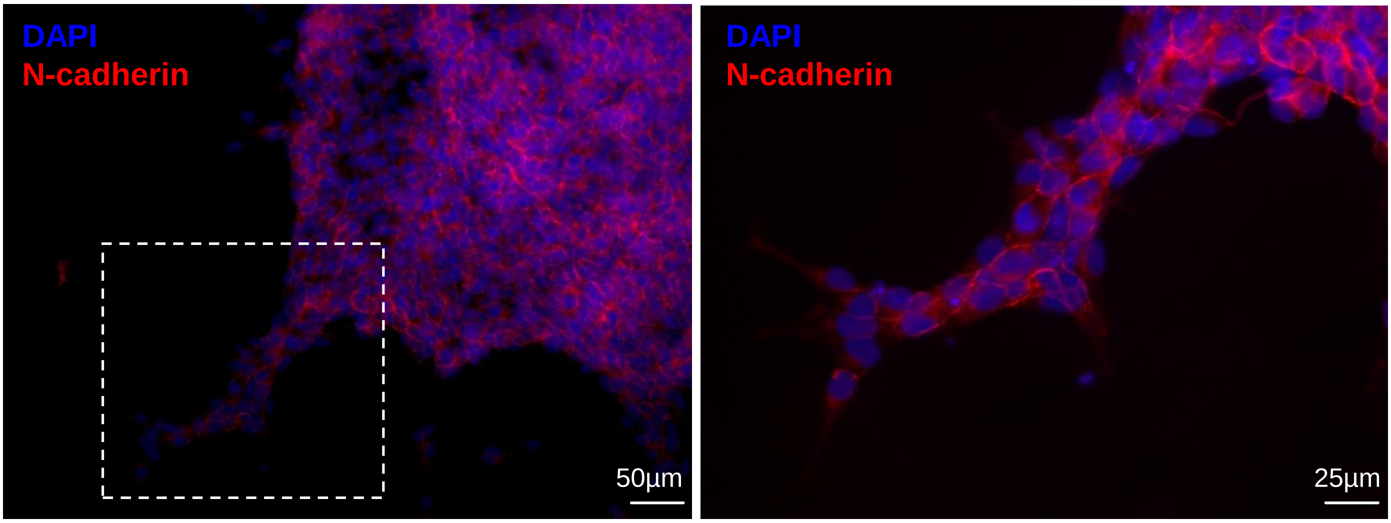

Supplemental Figure 3

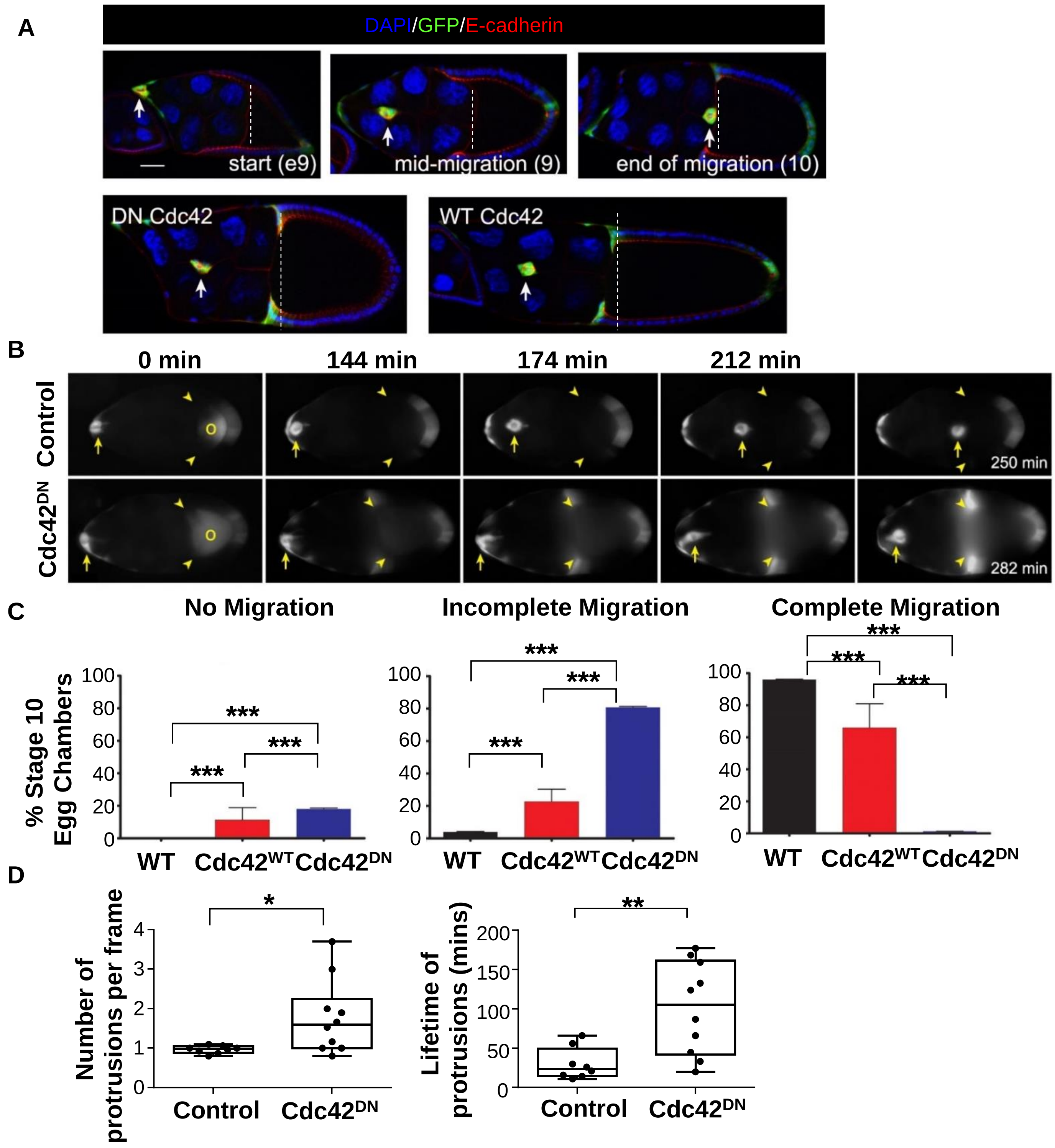

**Supplemental Figure 4**

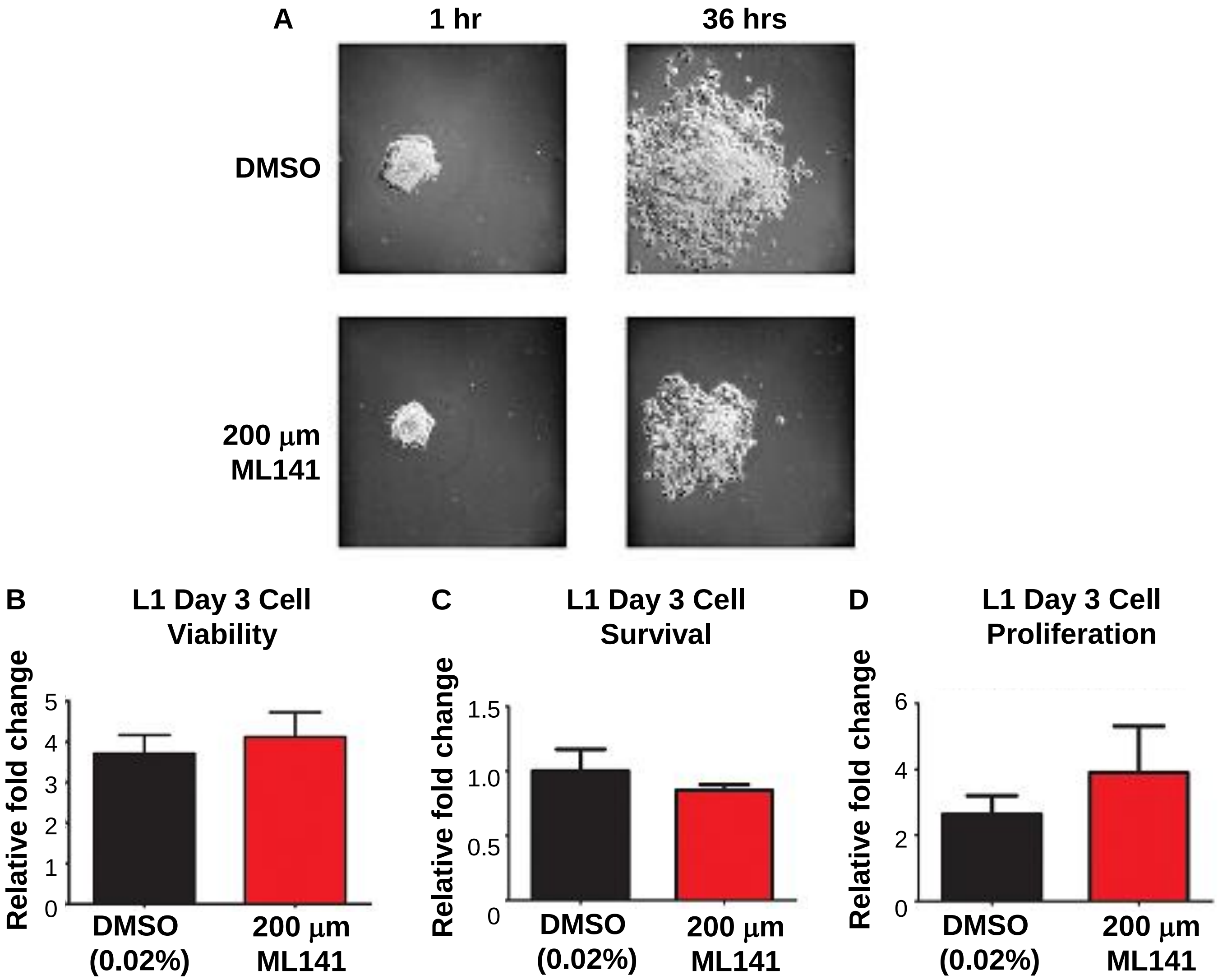

| Group | Gene Name (synonym) | Gene Symbo | BC migration defects | Rationale for inclusion | Human homolog(s) | Human homolog symbol(s) | Dataset | p value | Exp. = increased or decreased surviva | Log rank test statistic | Reference for BC migration |
|---|---|---|---|---|---|---|---|---|---|---|---|
| **1** | bazooka | baz | Yes | PDZ domain gene | PARD3 | PAR3 | GBMLGG | 0.000 | increased | 91.21 | Pinheiro and Montell, 2004 |
| **Multiple RNAi lines with BC phenotypes** | big bang | bbg | Yes | PDZ domain gene | *none* | none | ND | | | | Aranjuez et al., 2012 |
| | CASK ortholog | CASK | Yes | PDZ domain gene | Calcium/calmodulin dependent serine-protein kinase | CASK | GBMLGG | 0.000 | increased | 56.46 | Aranjuez et al., 2012 |
| | CG5921 | CG5921 | Yes | PDZ domain gene | USH1 protein network component harmonin | USH1C | GBMLGG | 0.000 | increased | 104.10 | Aranjuez et al., 2012 |
| | Discs large 5 | Dlg5 | Yes | PDZ domain gene | discs large MAGUK scaffold protein 5 | DLG5 | GBMLGG | 0.000 | increased | 120.80 | Aranjuez et al., 2012 |
| | Drop out | Dop | Yes | PDZ domain gene | microtubule associated serine/threonine kinase 1 (MAST1) | MAST1 | GBMLGG | 0.000 | increased | 77.81 | Aranjuez et al., 2012 |
| | *Drop out* | *Dop* | *Yes* | | MAST2 | MAST2 | GBMLGG | 0.001 | increased | 9.58 | |
| | *Drop out* | *Dop* | *Yes* | | MAST3 | MAST 3 | GBMLGG | 0.000 | increased | 24.17 | |
| | *Drop out* | *Dop* | *Yes* | | MAST4 | MAST4 | GBMLGG | 0.000 | increased | 13.71 | |
| | Lap1 | Lap1 | Yes | PDZ domain gene | erbb2 interacting protein | ERBB2 | GBMLGG | 0.000 | decreased | 44.97 | Aranjuez et al., 2012 |
| | *Lap1* | *Lap1* | *Yes* | | *leucine rich repeat containing 7* | *LRRC7* | GBMLGG | 0.000 | increased | 41.89 | |
| | LIM-kinase1 | LIMK1 | Yes | PDZ domain gene (actin regulator downstream of Rac, Rho, and Cdc42 GTPases) | LIM domain kinase 1, LIMK2 | LIMK1, LIMK2 | GBMLGG | 0.000 | decreased | 73.62 | Aranjuez et al., 2012 |
| | par-6 | par-6 | Yes | PDZ domain gene | par-6 family cell polarity regulator alpha | PARD6A (Par6) | GBMLGG | 0.000 | increased | 35.10 | Pinheiro and Montell, 2004 |
| | *par-6* | *par-6* | *Yes* | | *par-6 family cell polarity regulator beta* | *PARD6B (Par6)* | GBMLGG | 0.006 | decreased | 7.501 | |
| | *par-6* | *par-6* | *Yes* | | *par-6 family cell polarity regulator gamma* | *PARD6G (Par6)* | GBMLGG | 0.000 | decreased | 27.60 | |
| | PatJ | Patj | Yes | PDZ domain gene | multiple PDZ domain crumbs cell polarity complex component | MPDZ | GBMLGG | 0.000 | increased | 95.82 | Aranjuez et al., 2012; Wang et al., 2018 |
| | *PatJ* | *Patj* | *Yes* | | *PATJ, crumbs cell polarity complex component*[3] | *PATJ* | GBMLGG | | | | |
| | PDZ-GEF (Dizzy) | PDZ-GEF | Yes | PDZ domain gene (GEF for Rap1GTPase) | Rap guanine nucleotide exchange factor 2 (PDZ-GEF1) | RAPGEF2 | GBMLGG | 0.000 | increased | 133.8 | Aranjuez et al., 2012; Sawant et al., 2018 |
| | *PDZ-GEF (Dizzy)* | *PDZ-GEF* | *Yes* | | *Rap guanine nucleotide exchange factor 6 (PDZ-GEF2)* | *RAPGEF6* | GBMLGG | | | | |
| | Rab3 interacting molecule | Rim | Yes | PDZ domain gene | regulating synaptic membrane exocytosis 1 | RIMS1 | GBMLGG | 0.000 | increased | 39.23 | Aranjuez et al., 2012 |
| | *Rab3 interacting molecule* | *Rim* | *Yes* | | regulating synaptic membrane exocytosis 2 | RIMS2 | GBMLGG | 0.000 | increased | 74.09 | |
| | stardust | sdt | Yes | PDZ domain gene | membrane palmitoylated protein 5 | MPP5 | GBMLGG | 0.446 | none | 0.58 | Aranjuez et al., 2012; Wang et al., 2018 |
| | veli | veli | Yes | PDZ domain gene | lin-7 homolog A, crumbs cell polarity complex component (LIN7A) | LIN7A | GBMLGG | 0.000 | decreased | 15.87 | Aranjuez et al., 2012 |
| | *veli* | *veli* | *Yes* | | LIN7B | LIN7B | GBMLGG | 0.000 | none | 0.53 | |
| | *veli* | *veli* | *Yes* | | LIN7C | LIN7C | GBMLGG | 0.10 | none | 2.57 | |
| **2** | CG34375 | CG34375 | Yes | PDZ domain gene | *none* | none | ND | | | | Aranjuez et al., 2012 |
| **Single RNAi line with BC phenotypes** | CG42319 | CG42319 | Yes | PDZ domain gene | *none* | none | ND | | | | Aranjuez et al., 2012 |
| | CG42788 | CG42788 | Yes | PDZ domain gene | FERM and PDZ domain containing 4 (FRMPD4) | FRMPD4 | GBMLGG | 0.000 | increased | 22.96 | Aranjuez et al., 2012 |
| | *CG42788* | *CG42788* | *Yes* | | FRMPD1 | FRMPD1 | GBMLGG | 0.000 | increased | 111.9 | |
| | *CG42788* | *CG42788* | *Yes* | | FRMPD3 | FRMPD3 | GBMLGG | | | | |
| | CG6688 | CG6688 | Yes | PDZ domain gene | *none* | none | ND | | | | Aranjuez et al., 2012 |
| | Dishevelled | dsh | Yes | PDZ domain gene | dishevelled segment polarity protein 1 (DVL1), DVL2, DVL3 | DVL1 | GBMLGG | 0.000 | increased | 25.38 | Aranjuez et al., 2012; Bastock and Strutt, 2007 |
| | *Dishevelled* | *dsh* | *Yes* | | | DVL2 | GBMLGG | 0.000 | increased | 10.55 | |
| | *Dishevelled* | *dsh* | *Yes* | | | DVL3 | GBMLGG | 0.000 | increased | 14.23 | |
| | Exchange factor for Arf 6 | Efa6 | Yes | PDZ domain gene | pleckstrin and Sec7 domain containing (PSD) | PSD | GBMLGG | 0.000 | increased | 97.86 | Aranjuez et al., 2012 |
| | *Exchange factor for Arf 6* | *Efa6* | *Yes* | | PSD2 | PSD2 | GBMLGG | 0.000 | increased | 30.44 | |
| | *Exchange factor for Arf 6* | *Efa6* | *Yes* | | PSD3 | PSD3 | GBMLGG | 0.000 | increased | 20.03 | |
| | *Exchange factor for Arf 6* | *Efa6* | *Yes* | | PSD4 | PSD4 | GBMLGG | 0.000 | decreased | 31.76 | |
| | HTRA2-related serine protease | HtrA2 | Yes | PDZ domain gene | HtrA serine peptidase 1 (HTRA1) | HTRA1 | GBMLGG | 0.612 | none | 0.2571 | Aranjuez et al., 2012 |
| | *HTRA2-related serine protease* | *HtrA2* | *Yes* | | HTRA2 | HTRA2 | GBMLGG | 0.708 | none | 0.1398 | |
| | *HTRA2-related serine protease* | *HtrA2* | *Yes* | | HTRA3 | HTRA3 | GBMLGG | 0.000 | decreased | 147.9 | |
| | *HTRA2-related serine protease* | *HtrA2* | *Yes* | | HTRA4 | HTRA4 | GBMLGG | 0.000 | decreased | 48.08 | |

| Group | Gene Name (synonym) | Gene Symbol | BC migration defects | Rationale for inclusion | Human homolog(s) | Human homolog symbol(s) | Dataset | p value | Exp. = increased or decreased survival | Log rank test statistic | Reference for BC migration |
|---|---|---|---|---|---|---|---|---|---|---|---|
| | Magi | Magi | Yes | PDZ domain gene | membrane associated guanylate kinase, WW and PDZ domain containing 1 (MAGI1) | MAGI1 | GBMLGG | 0.000 | increased | 21.79 | Aranjuez et al., 2012 |
| | *Magi* | *Magi* | *Yes* | | MAGI2 | MAGI2 | GBMLGG | 0.5615 | none | 0.3371 | |
| | *Magi* | *Magi* | *Yes* | | MAGI3 | MAGI3 | GBMLGG | 0.000 | increased | 14.80 | |
| | *Magi* | *Magi* | *Yes* | | MAGI4 | MAGI4 | GBMLGG | | | | |
| | menage a trois | metro | Yes | PDZ domain gene | membrane palmitoylated protein 7 | MPP7 | GBMLGG | 0.000 | decreased | 61.31 | Aranjuez et al., 2012 |
| | polychaetoid | pyd | Yes | PDZ domain gene | tight junction protein 1 (TJP1) | TJP1 | | | | | Aranjuez et al., 2012 |
| | *polychaetoid* | *pyd* | *Yes* | | TJP2 | TJP2 | GBMLGG | 0.000 | increased | 127.5 | |
| | Ptpmeg | Ptpmeg | Yes | PDZ domain gene | protein tyrosine phosphatase, non-receptor type 4 | PTPN4 | GBMLGG | 0.000 | increased | 28.60 | Aranjuez et al., 2012 |
| | Rho GTPase activating protein at 19D | RhoGAP19D | Yes | PDZ domain gene (GAP for Rho GTPase) | Rho GTPase activating protein 21 (ARHGAP21) | ARHGAP21 | GBMLGG | 0.000 | increased | 49.85 | Aranjuez et al., 2012 |
| | *Rho GTPase activating protein at 19D* | *RhoGAP19D* | *Yes* | | ARHGAP23 | ARHGAP23 | GBMLGG | | | | |
| | Rhophilin | Rhp | Yes | PDZ domain gene | rhophilin Rho GTPase binding protein 1 (RHPN1) | RHPN1 | GBMLGG | 0.043 | increased | 4.095 | Aranjuez et al., 2012 |
| | *Rhophilin* | *Rhp* | *Yes* | | RHPN2 | RHPN2 | GBMLGG | 0.000 | decreased | 45.80 | |
| | Slip1 | Slip1 | Yes | PDZ domain gene | PDZ domain containing 4 | PDZD4 | GBMLGG | 0.000 | increased | 108.3 | Aranjuez et al., 2012 |
| | *Slip1* | *Slip1* | *Yes* | | *PDZ domain containing ring finger 3 (PDZRN3)* | PDZRN3 | GBMLGG | 0.002 | increased | 9.03 | |
| | *Slip1* | *Slip1* | *Yes* | | *PDZRN4* | *PDZRN4* | GBMLGG | 0.000 | increased | 25.11 | |
| | Syntrophin-like 2 | Syn2 | Yes | PDZ domain gene | syntrophin gamma 1 (SNTG1) | *SNTG1* | GBMLGG | 0.000 | increased | 16.50 | Aranjuez et al., 2012 |
| | *Syntrophin-like 2* | *Syn2* | *Yes* | | SNTG2 | SNTG2 | GBMLGG | 0.002 | decreased | 9.112 | |
| | Z band alternatively spliced PDZ-motif protein 52 | Zasp52 | Yes | PDZ domain gene | PDZ and LIM domain 7 (PDLIM7) | PDLIM7 | GBMLGG | 0.002 | decreased | 124 | Aranjuez et al., 2012 |
| | Z band alternatively spliced PDZ-motif protein 67 | Zasp67 (CG14168) | Yes | PDZ domain gene | *none* | none | ND | | | | Aranjuez et al., 2012 |
| 3 | atypical Protein Kinase C | aPKC | Yes | Binds to Par-6 | protein kinase C zeta | PRKCζ | GBMLGG | 0.000 | increased | 46.75 | Pinheiro and Montell, 2004; Wang et al., 2018 |
| **Related genes (positive role in BC migration and/or known PDZ-gene interactors)** | *atypical Protein Kinase C* | *aPKC* | *Yes* | | *protein kinase C iota* | *PRKCι* | GBMLGG | | | | |
| | Cdc42 | Cdc42 | Yes | Small GTPase, binds to Par-6 | CDC42 | CDC42 | GBMLGG | 0.000 | decreased | 98.90 | Llense and Martin-Blanco, 2008; Colombié et al., 2017; this study |
| | Cin85 and CD2AP related | Cindr | Yes | Binds to Efa6, required for BC migration | SH3 domain containing kinase binding protein 1 | SH3KBP1 | GBMLGG | 0.000 | decreased | 33.21 | Quinones et al., 2010 |
| | *Cin85 and CD2AP related* | *Cindr* | *Yes* | | *CD2 associated protein* | *CD2AP* | GBMLGG | 0.00 | decreased | 33.61 | |
| | Rac1 | Rac1 | Yes | Small GTPase, LIMK1 is downstream, Sif/TIAM1 is a GEF, required for BC migration | Rac family small GTPase 1 (RAC1) | RAC1 | GBMLGG | 0.000 | decreased | 31.82 | Murphy and Montell, 1996; Wang et al., 2010 |
| | Rac 2 | Rac 2 | Yes | | RAC2 | RAC2 | GBMLGG | 0.000 | decreased | 98.00 | |
| | *Rac1, Rac 2* | *Rac1, Rac 2* | *Yes* | | RAC3 | RAC3 | GBMLGG | 0.000 | increased | 52.02 | |
| | Rap1 | Rap1 | Yes | Small GTPase, PDZ-GEF is GEF for Rap1 | RAP1A | RAP1A | GBMLGG | 0.000 | decreased | 78.03 | Chang et al., 2018; Sawant et al., 2018 |
| | *Rap1* | *Rap1* | *Yes* | | RAP1B | RAP1B | GBMLGG | 0.000 | decreased | 133 | |
| | Shotgun | Shg | Yes | Baz binds to Shg, required for BC migration | E-cadherin | CDH1 | GBMLGG | 0.484 | none | 0.488 | Niewiadomska et al., 1999 |
| | Spaghetti squash | Sqh | Yes | Baz binds to Sqh, required for BC migration | myosin light chain 12A | MYL12A | GBMLGG | 0.000 | decreased | 138.00 | Fulga and Rørth, 2003; Majumder et al., 2012 |
| | *Spaghetti squash* | *Sqh* | *Yes* | | myosin light chain 9 | MYL9 | GBMLGG | 0.000 | decreased | 63.36 | |
| | α-catenin | α-cat | Yes | Baz binds to α-cat, required for BC migration | catenin alpha 1 (CTNNA1) | CTNNA1 | GBMLGG | 0.000 | decreased | 110.00 | Sarpal et al., 2012 |
| | *α-catenin* | *α-cat* | *Yes* | | CTNNA2 | CTNNA2 | GBMLGG | 0.000 | increased | 11.96 | |
| | *α-catenin* | *α-cat* | *Yes* | | CTNNA3 | CTNNA3 | GBMLGG | 0.000 | increased | 67.51 | |
| **4** | canoe | cno | No | PDZ domain gene | afadin, adherens junction formation factor | AFDN | ND | | | | Aranjuez et al., 2012 |
| **No phenotypes in BCs by RNAi[1]** | CG10362 | CG10362 | No | PDZ domain gene | PDZ domain containing 8 | PDZD8 | ND | | | | Aranjuez et al., 2012 |
| | CG15617 | CG15617 | No | PDZ domain gene | *none* | none | ND | | | | Aranjuez et al., 2012 |

| Group | Gene Name (synonym) | Gene Symbol | BC migration defects | Rationale for inclusion | Human homolog(s) | Human homolog symbol(s) | Dataset | p value | Exp. = increased or decreased survival | Log rank test statistic | Reference for BC migration |
|---|---|---|---|---|---|---|---|---|---|---|---|
| | CG15803 | CG15803 | No | PDZ domain gene | *none* | none | ND | | | | Aranjuez et al., 2012 |
| | CG32758 | CG32758 | No | PDZ domain gene | sorting nexin family member 27 | SNX27 | ND | | | | Aranjuez et al., 2012 |
| | CG3402 | CG3402 | No | PDZ domain gene | Tax1 binding protein 3 | TAX1BP3 | ND | | | | Aranjuez et al., 2012 |
| | CG43707 | CG43707 | No | PDZ domain gene | *none* | none | ND | | | | Aranjuez et al., 2012 |
| | CG9588 | CG9588 | No | PDZ domain gene | proteasome 26S subunit, non-ATPase 9 | PSMD9 | ND | | | | Aranjuez et al., 2012 |
| | connector enhancer of ksr | cnk | No | PDZ domain gene | connector enhancer of kinase suppressor of Ras 2 (CNKSR2), CNKSR1, CNKSR3 | CNKSR2, CNKSR1, CNKSR3 | ND | | | | Aranjuez et al., 2012 |
| | discs large 1 | dlg1 | No | PDZ domain gene | discs large MAGUK scaffold protein 1 (DLG1), DLG2, DLG3, DLG4 | DLG1, DLG2, DLG3, DLG4 | ND | | | | Aranjuez et al., 2012 |
| | dyschronic | dysc | No | PDZ domain gene | whirlin | WHRN | ND | | | | Aranjuez et al., 2012 |
| | Fife | Fife | No | PDZ domain gene | *none* | none | ND | | | | Aranjuez et al., 2012 |
| | Glutamine receptor binding protein | Grip | No | PDZ domain gene | glutamate receptor interacting protein 1 (GRIP1), GRIP2 | GRIP1, GRIP2 | ND | | | | Aranjuez et al., 2012 |
| | Grasp65 | Grasp65 | No | PDZ domain gene | golgi reassembly stacking protein 1 (GORASP1), GORASP2 | GORASP1, GORASP2 | ND | | | | Aranjuez et al., 2012 |
| | inactivation no afterpotential D | inaD | No | PDZ domain gene | *none* | none | ND | | | | Aranjuez et al., 2012 |
| | kermit | kermit | No | PDZ domain gene | GIPC PDZ domain containing family member 1 (GIPC1), GIPC2, GIPC3 | GIPC1, GIPC2, GIPC3 | ND | | | | Aranjuez et al., 2012 |
| | locomotion defects | loco | No | PDZ domain gene | regulator of G protein signaling 12 | RGS12 | ND | | | | Aranjuez et al., 2012 |
| | PICK1 | PICK1 | No | PDZ domain gene | protein interacting with PRKCA 1 | PICK1 | ND | | | | Aranjuez et al., 2012 |
| | Prosap | Prosap | No | PDZ domain gene | SH3 and multiple ankyrin repeat domains 1 (SHANK1), SHANK2, SHANK3 | SHANK1, SHANK2, SHANK3 | ND | | | | Aranjuez et al., 2012 |
| | Protostome-specific GEF | PsGEF | No | PDZ domain gene | *none* | none | ND | | | | Aranjuez et al., 2012 |
| | RhoGAP100F | RhoGAP100F | No | PDZ domain gene | synapse defective Rho GTPase homolog 1 (SYDE1), SYDE2 | SYDE1, SYDE2 | ND | | | | Aranjuez et al., 2012 |
| | RhoGEF2 | RhoGEF2 | No | PDZ domain gene | Rho guanine nucleotide exchange factor 1 (ARHGEF1) | ARHGEF1 | ND | | | | Aranjuez et al., 2012 |
| | *RhoGEF2* | *RhoGEF2* | *No* | | Rho guanine nucleotide exchange factor 11 | ARHGEF11 | ND | | | | |
| | *RhoGEF2* | *RhoGEF2* | *No* | | Rho guanine nucleotide exchange factor 12 | ARHGEF12 | ND | | | | |
| | scribbled | scrib | No | PDZ domain gene | leucine rich repeat containing 1 | LRRC1 | ND | | | | Aranjuez et al., 2012 |
| | short spindle 6 (CG6619) | ssp6 | No | PDZ domain gene | *none* | none | ND | | | | Aranjuez et al., 2012 |
| | Spinophilin | Spn | No | PDZ domain gene | protein phosphatase 1 regulatory subunit 9A (PPP1R9A), PPP1R9B | PPP1R9A, PPP1R9B | ND | | | | Aranjuez et al., 2012 |
| | sprite | sprt | No | PDZ domain gene | *none* | none | ND | | | | Aranjuez et al., 2012 |
| | still life | sif | No [1], Yes [2] | PDZ domain gene (GEF for Rac GTPase) | T-cell lymphoma invasion and metastasis 1 | TIAM1 | ND | | | | [1] Aranjuez et al., 2012 no phenotype; [2] Wang et al. 2018 required for BC migration |
| | Syntrophin-like 1 | Syn1 | No | PDZ domain gene | syntrophin beta 1 (SNTB1), SNTB2 | SNTB1, SNTB2 | ND | | | | Aranjuez et al., 2012 |
| | varicose | vari | No | PDZ domain gene | membrane palmitoylated protein 2 | MPP2 | ND | | | | Aranjuez et al., 2012 |
| | *varicose* | *vari* | *No* | | membrane palmitoylated protein 6 | MPP6 | ND | | | | |
| | X11L | X11L | No | PDZ domain gene | amyloid beta precursor protein binding family A member 1 (APBA1), APBA2, APBA3 | APBA1, APBA2, APBA3 | ND | | | | Aranjuez et al., 2012 |
| | X11Lbeta | X11Lbeta | No | PDZ domain gene | amyloid beta precursor protein binding family A member 1 (APBA1), APBA2, APBA3 | APBA1, APBA2, APBA3 | ND | | | | Aranjuez et al., 2012 |
| | Z band alternativel spliced PDZ-motif protein 66 | Zasp66 | No | PDZ domain gene | LIM domain binding 3 | LDB3 | ND | | | | Aranjuez et al., 2012 |
| **5** | arc | a | **I** (ND) | PDZ domain gene | PDZ domain containing 2 | PDZD2 | ND | | | | Aranjuez et al., 2012 |
| **Not tested in BCs[2]** | Myosin heavy chain-like | Mhcl | **I** (ND) | PDZ domain gene | myosin XVIIIA | MYO18A | ND | | | | Aranjuez et al., 2012 |

[1]No phenotype in BCs, at least in primary RNAi screen (Aranjuez et al., 2012); *sif* has subsequently been shown to be required for border cell migration (Wang et al., 2018).

[2]No specific RNAi line was available at the time of the primary RNAi screen (Aranjuez et al., 2012).

[3]Genes in red were not present in the GBMLGG database.
